# Highly efficient direct conversion of human monocytes into neuronal cells using a small molecule combination

**DOI:** 10.1101/803254

**Authors:** Itaru Ninomiya, Masato Kanazawa, Akihide Koyama, Masahiro Hatakeyama, Osamu Onodera

**Affiliations:** Department of Neurology, Brain Research Institute, Niigata University, 1-757 Asahimachi-dori, Chuoku, Niigata 951-8585, Japan

**Author notes:** **Author Contributions** Conceptualization, I.N.; Methodology, I.N., M.K., and A.K.; Investigation, I.N., and A.K.; Writing – Original Draft, I.N.; Writing – Review & Editing, I.N., M.K., A.K., M.H., and O.O.; Resources, M.K., and O.O.; Supervision, O.O.

**Keywords:** induced neuronal cells, direct reprogramming, transdifferentiation, monocytes, iN cells, direct conversion, small molecules

## Abstract

Previous studies reported that human fibroblasts and astrocytes were successfully converted into neuronal cells by small molecules without introducing ectopic transgenes. Induced neuronal cells—reprogrammed directly from dermal fibroblasts or brain astrocytes—were obtained from some donors; however, the clinical applications of this approach would be limited because it requires an invasive biopsy to harvest enough cells for derivation. Here, we report that adult human peripheral blood monocytes may be directly converted into neuron-like cells using only a combination of small molecules without transgene integration. This method enables neuronal cell generation from TUJ1-positive cells after 3 days of induction (at over 80% conversion efficacy). These cells presented neuronal morphologies and markers, suggesting that terminally differentiated human cells may be efficiently transdifferentiated into a distantly related lineage. Overall, our study provides a strategy to develop neuronal cells directly from human adult peripheral blood monocytes using a generate transgene-free, chemical-only approach.

## Introduction

One of the most promising therapies for neurological diseases such as: Parkinson’s disease, Alzheimer’s disease, stroke, and traumatic brain injury, is neuronal replacement using exogenous cells. Because the regenerative capacity of adult brains is limited (Goldman et al., 2016), various attempts have been made in the field of regenerative medicine to develop exogenous neurons. One way to create such neurons is to reprogram patients’ terminally differentiated somatic cells into induced pluripotent stem (iPS) cells via forcing the expression of a specific set of transcription factors (TFs). iPS-derived neuronal cells are promising foundational resources for the restoration of neuronal function (Lindvall and Kokaia, 2006). However, because the pluripotency of iPS cells may vary from cell to cell, the differentiation capacity of iPS cells likewise varies (Lindvall and Kokaia, 2006; Liu et al., 2013; Nichols et al., 2009). Moreover, risk of tumorigenesis, and the difficulty associated with delivering iPS-derived neurons into the brain, are barriers to their clinical application. Therefore, a safe and efficient method of exogenous neuronal cell generation is essential for their use in clinical applications.

TFs are considered the major determinants of specific cell lineages and conversions (Mertens et al., 2016; Xu et al., 2015). Small molecules improve TFs-mediated neuronal conversion efficiency (Gascón et al., 2016; Ladewig et al., 2012; Liu et al. 2013). Interestingly, some reports have also revealed recently that specific combinations of small molecules, without integration of ectopic transgenes, can activate key neuronal TFs in fibroblasts and astrocytes—reprogramming them into neuronal cells (Cheng et al., 2015; Hu et al., 2015; Li et al., 2015; Zhang et al., 2015; Gao et al., 2017). Importantly, in these conversion methods, the exogenous cells do not pass through a pluripotent intermediate state, or require the integration of ectopic transgenes—preventing the risks of tumorigenesis or unexpected genomic gene modification. However, because an invasive biopsy is often necessary to obtain fibroblasts and astrocytes, it is difficult to repetitively harvest these cell sources and acquire sufficient cell numbers for clinical usefulness. Therefore, these cells must be obtained less invasively for these methods to be considered for clinical application.

Human blood cells are ideal cell sources for transdifferentiation because they are easy to harvest via simple venipuncture. Peripheral blood mononuclear cells (PBMCs) contain a population of monocytes, lymphocytes, mesenchymal progenitor cells and stem cells. Tanabe et al. reported that human peripheral blood T cells may be transdifferentiated into neuron-like cells by small molecules (Tanabe et al., 2018). However, the conversion method they described requires the integration of ectopic transgenes. Herein we report that neuron-like cells may be directly induced from adult peripheral monocytes using a combination of small molecules without integrating ectopic transgenes.

## Results

### Direct conversion of human peripheral blood mononuclear cells into neuronal cells

To investigate whether PBMCs could be converted into neurons, we prepared human PBMCs from fresh human adult peripheral blood using gradient centrifugation. The PBMCs were cultured in X-VIVO15 medium for 2 hours. The culture medium—containing floating cells—was then discarded. Remaining adherent cells on the culture plate were incubated with several combinations of small molecules for 3 days (Figure 1A). We found that 3 day incubation with the combination of small molecules CDFRVY (CHIR99021, Dorsomorphin, Forskolin, Repsox, Valproic acid, and Y27632) drastically changed the PBMC cell morphology—which presented neurite-like structures after treatment. These induced neuronal (iN) cells expressed TUJ1, a known neuronal marker (Figures 1B, right panel; and C). In contrast, no significant morphological change was observed in the control group—where no small molecules were added (Figure 1B, left panel). Because PBMCs consist of several cell types—such as monocytes and lymphocytes—we further investigated which PBMC cell type was converted into iN cells. Using magnetic beads conjugated with anti-CD14 antibodies, PBMCs were separated into monocytes and other cells (monocyte-depleted PBMCs) which mostly consisted of lymphocytes. The monocytes and monocyte-depleted PBMCs were cultured separately with CDFRVY small molecules for 3 days (Figure 1D). We found that only monocytes were converted into neuronal cells and no significant morphological change was observed in the monocyte-depleted PBMCs after 3 days of culture (Figures 1E and 1F). Because the monocyte-depleted PBMCs consisted of mainly lymphocytes, which rarely adhere on culture plate, very few cells were observed on day 3. These findings confirmed that the neuronal cells were converted from monocytes.

**Figure 1.**
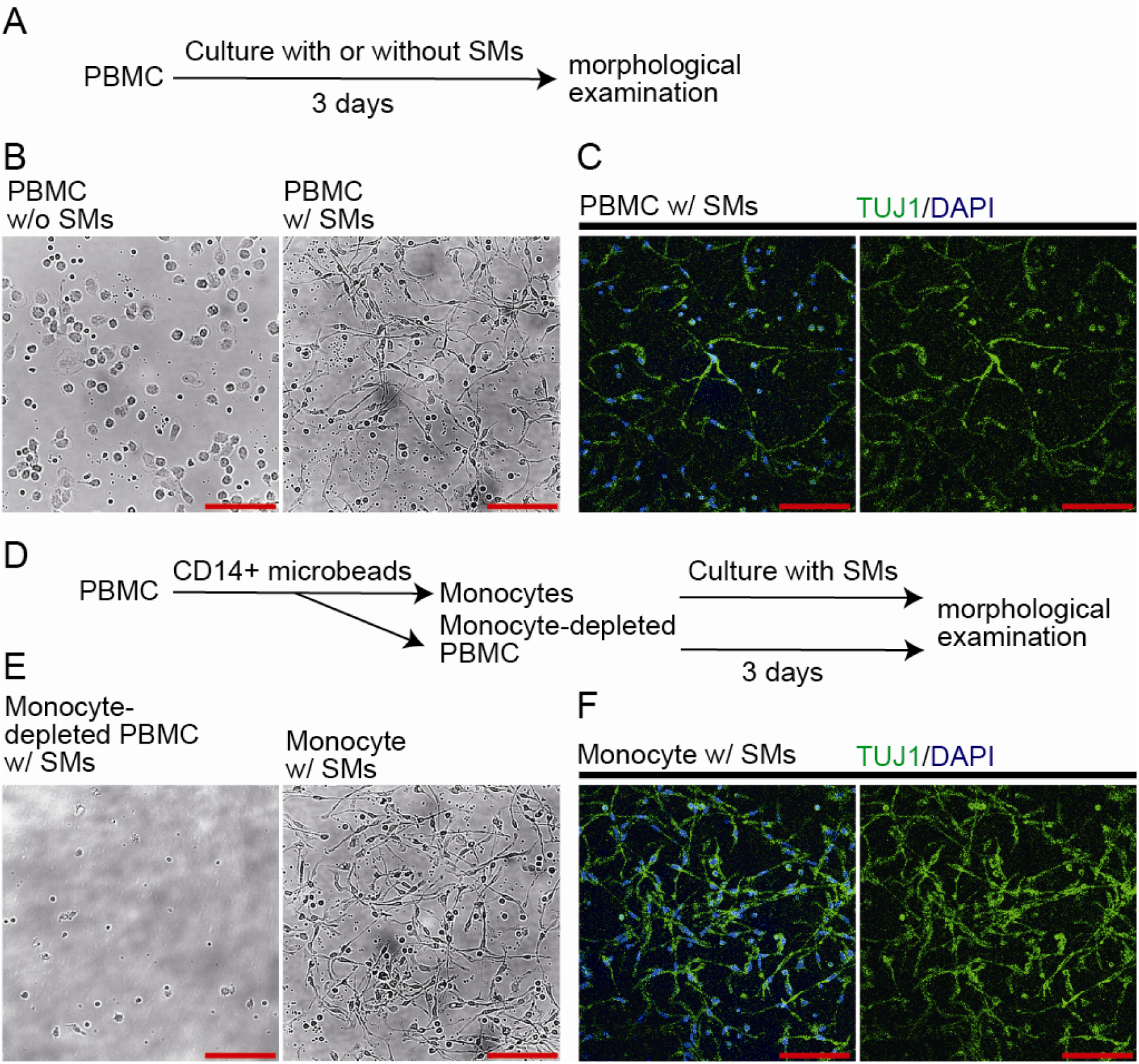
Direct conversion of human peripheral blood mononuclear cells (PBMCs) into neuronal cells. (A) Scheme representing of the small-molecule treatment of PBMCs. SMs: small molecules (B) Phase contrast images of non-CDFRVY-treated PBMC (left) and CDFRVY treated PBMC (right). Scale bars, 100 µm. (C) TUJ1-positive cells. Induced from PBMCs by CDFRVY small molecules (D) Diagram of monocyte and monocyte-depleted PBMC isolation. Monocytes and monocyte-depleted PBMCs were separately cultured with small molecules for 3 days. (E) Phase contrast images of CDFRVY treated monocyte-depleted PBMC (left) and CDFRVY treated monocytes (right). Scale bars, 100 µm. (F) TUJ1-positive cells induced from monocytes by CDFRVY.

### CDFRVY-small-molecule treated human adult monocytes acquired neuronal properties

We investigated the expression of neural markers after small molecule induction (Figure 2A). Immunostaining revealed that the monocytic maker CD11b was detectable on day 0 but was not detected after CDFRVY treatment for 3 days (Figure 2B and C). TUJ1, DCX, and MAP2 were detectable at 3 days post induction (Figure 2D and E). These results suggested that adult monocytes acquired a neuronal fate after CDFRVY treatment. Based on TUJ1/DCX and TUJ1/MAP2 expression and cell morphology, the conversion efficiency was estimated to be 85.5 (±3.1) % and 7.4 (±0.9) %, respectively. The neuronal purity was estimated to be 91.4 (±1.3) % and 8.8 (±2.8) %, respectively (Figure 2F). Interestingly, monocytes could not be converted if even one of these small molecules was absent, indicating that the conversion required all of the CDFRVY small molecules.

**Figure 2.**
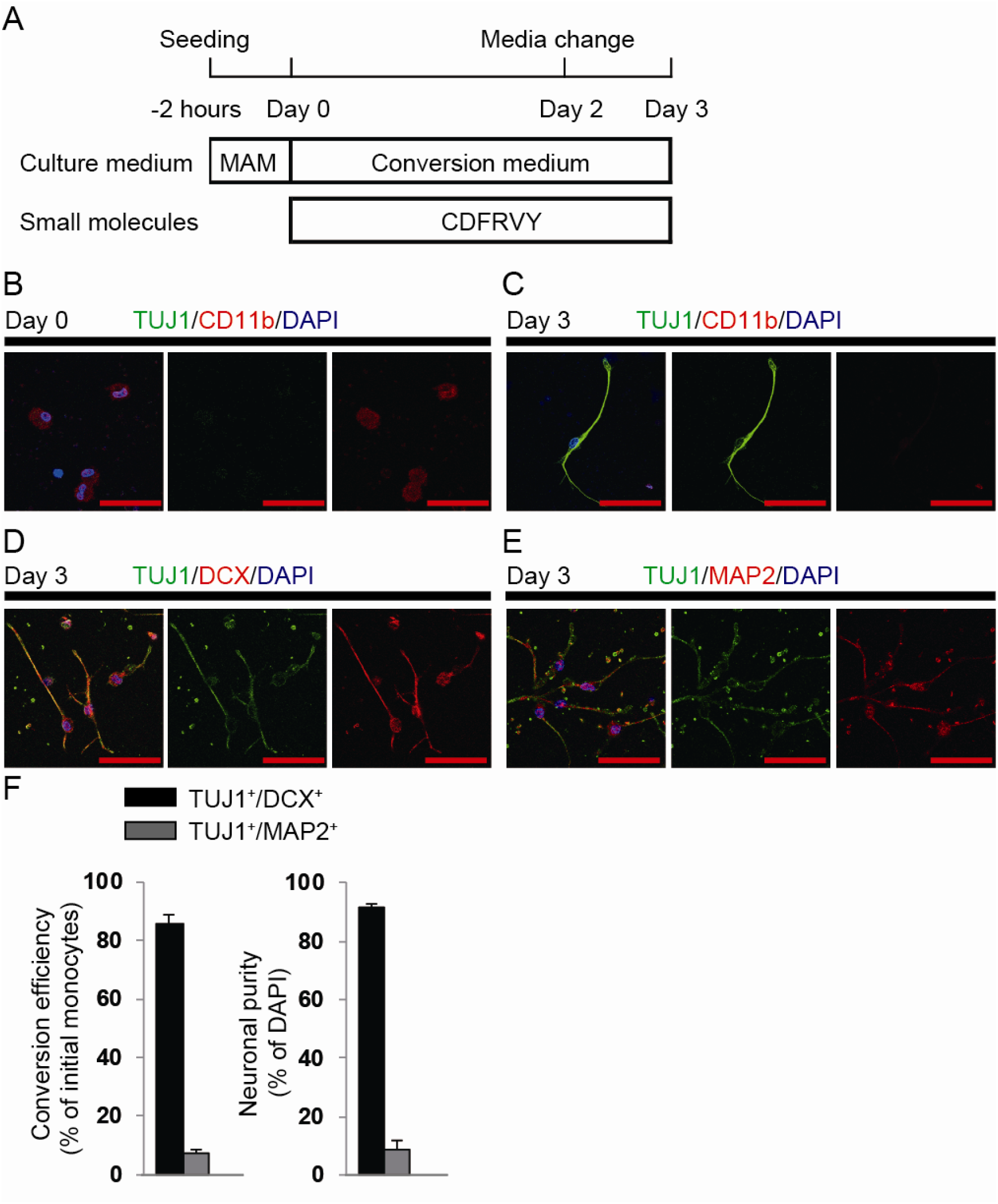
Induction of neuronal cells from human adult monocytes. (A) Schematic diagram showing the neuronal induction protocol. MAM, monocyte attachment medium; C, CHIR99021; D, Dorsomorphin; F, forskolin; R, Repsox; V, VPA; Y, Y27632. (B) Representative image of monocytes at day 0, showing round shape morphologies and CD11b (red) expression, but no TUJ1 (green) expression. Scale bar: 50 μm. (C-E) Induced neuronal cells display neuronal morphologies and express TUJ1 (green), DCX (red), and MAP2 (red) but not CD11b (red). Scale bar: 50 μm. (F) iN cell conversion efficiencies were presented as the percentage of induced TUJ1-positive and DCX-positive (or TUJ1-positive and MAP2-positive cells) on day 3 versus initial CD11b+ cells on day 0; neuronal purity was presented as the percentage of induced TUJ1-positive and DCX-positive, or TUJ1-positive and MAP2-positive cells on day 3 versus DAPI+ cells. (means ± SEM, n = 3 random selected ten fields)

### CDFRVY directly induced neuronal-master transcription factors and disrupted the monocyte-specific program during iNs generation

By absolute quantification analysis using droplet digital PCR (ddPCR), we found that the activation of neural-fate-determining genes and the downregulation of monocyte-fate-determining genes were induced within 1 day (Figures 3A). During the 3 days of chemical induction, ASCL1, BRN2, and NGN2 expression were initially upregulated and other neural-fate mastering genes were gradually induced. These results suggested that these genes may be involved in the first step of chemical reprogramming. Particularly, ASCL11, NEUROD1, and NGN2 are proneural genes that code a master transcription factor frequently reported (by various authors) to induce neuronal fate conversion (Matsuda et al., 2019; Tanabe et al., 2018; Li et al., 2015). By contrast, the monocyte and macrophage specific genes CD11b and Iba1 were gradually downregulated over 3 days. RT-qPCR analysis also revealed that neural-fate related genes—including proneural genes—were gradually upregulated over 3 days by CDFRVY treatment (Figure 3B). Western blot analysis revealed that TUJ1, DCX, and ASCL1 were increasingly present on day 3 compared to day 0, while CD11b was less present on day 3 compared to day 0 (Figure 3C). These results suggested that CDFRVY treatment modified target cell fate through not only the activation of the target cell fate regulatory program, but also the repression of the original cell fate program.

**Figure 3.**
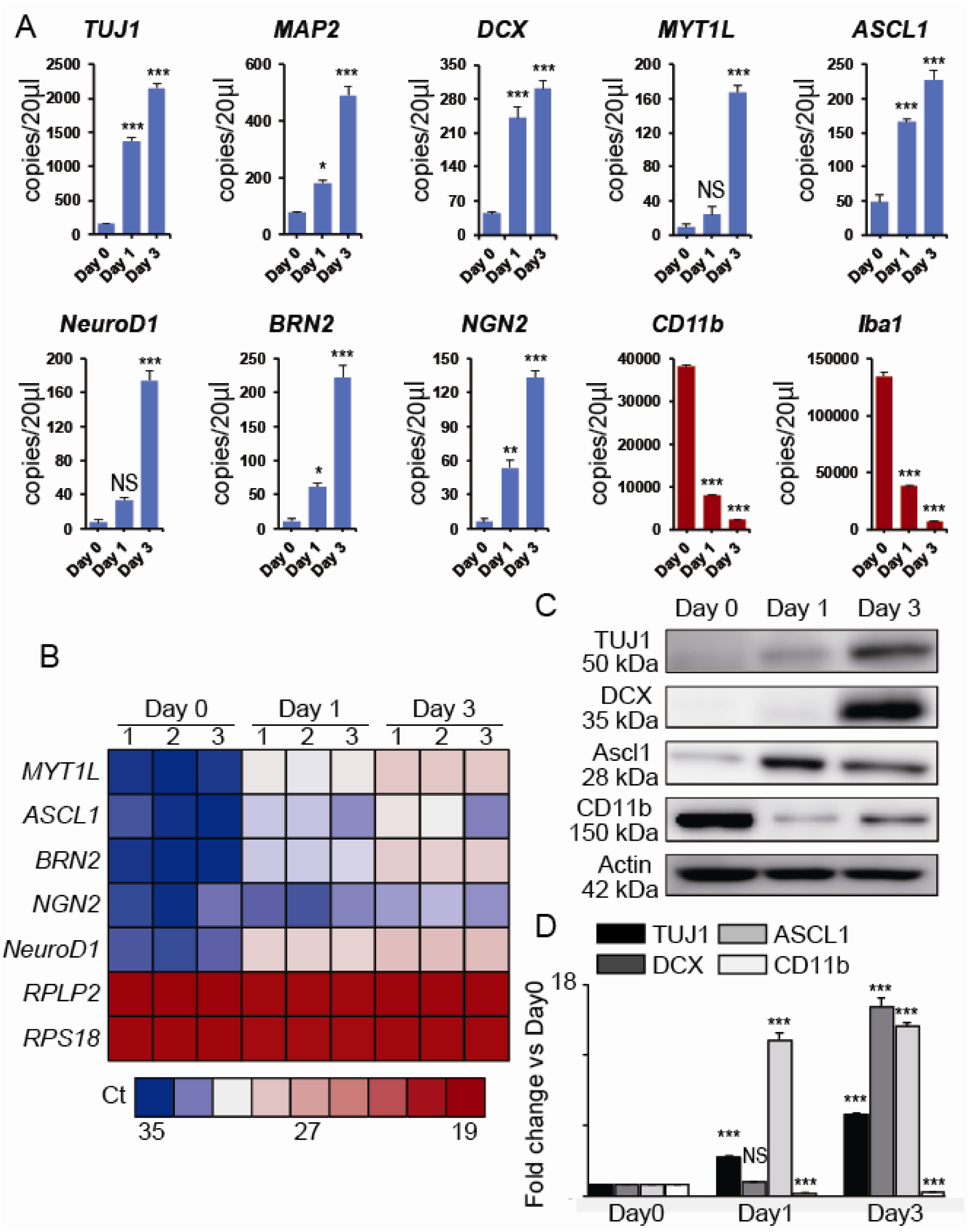
CDFRVY directly induced neuronal-master transcription factors and disrupted the monocyte-specific program during iNs generation. (A) mRNA copy number validation of monocyte- and neuron-enriched gene expression using absolute quantification analysis by droplet digital PCR. Monocyte-enriched genes are shown in red and neuron-enriched genes in blue. (mean ± SEM, n = 3 independent experiments, *p < 0.05; **p < 0.01; ***p < 0.001; NS, not significant; versus day 0 sample by unpaired t-test). (B) RT-qPCR analysis of the iN cells. Heat map showing expression change of neuronal genes over 3 days induction with CDFRVY. (C) Representative image of western blotting analysis of the iN cells. (D) Densitometry analysis for western blotting result. (mean ± SEM, n = 3 independent experiments, ***p < 0.001; NS, not significant; Fold change of TUJ1/actin, DCX/actin, ASCL1/actin, or CD11b/actin ratio versus day 0 sample, unpaired t-test).

### The blood-to-neuron conversion does not involve a proliferative neural progenitor state

Some groups have reported that blood cells can be converted into proliferative neural cells using subsets of iPS cell reprogramming factors (Matsui et al., 2012; Kim et al., 2011). In contrast, our reprogramming factors were not proliferative and our approach generated postmitotic neuronal cells directly. Although neuronal TFs such as NEUROD1, NGN2, and MYT1L were significantly upregulated between day 1 to day 3 (Figure 3A), neural progenitor cells (NPC) markers PAX6 and SOX2 remained at low expression levels and were not detectable by immunostaining between day 1 and 3 during reprogramming (Figures 4A-C). According to the results of EdU assay, cell proliferation was consistently undetectable during the conversion process (Figures 4D-E). Collectively, our results suggest that converted cells may not pass through the NPC stage.

**Figure 4.**
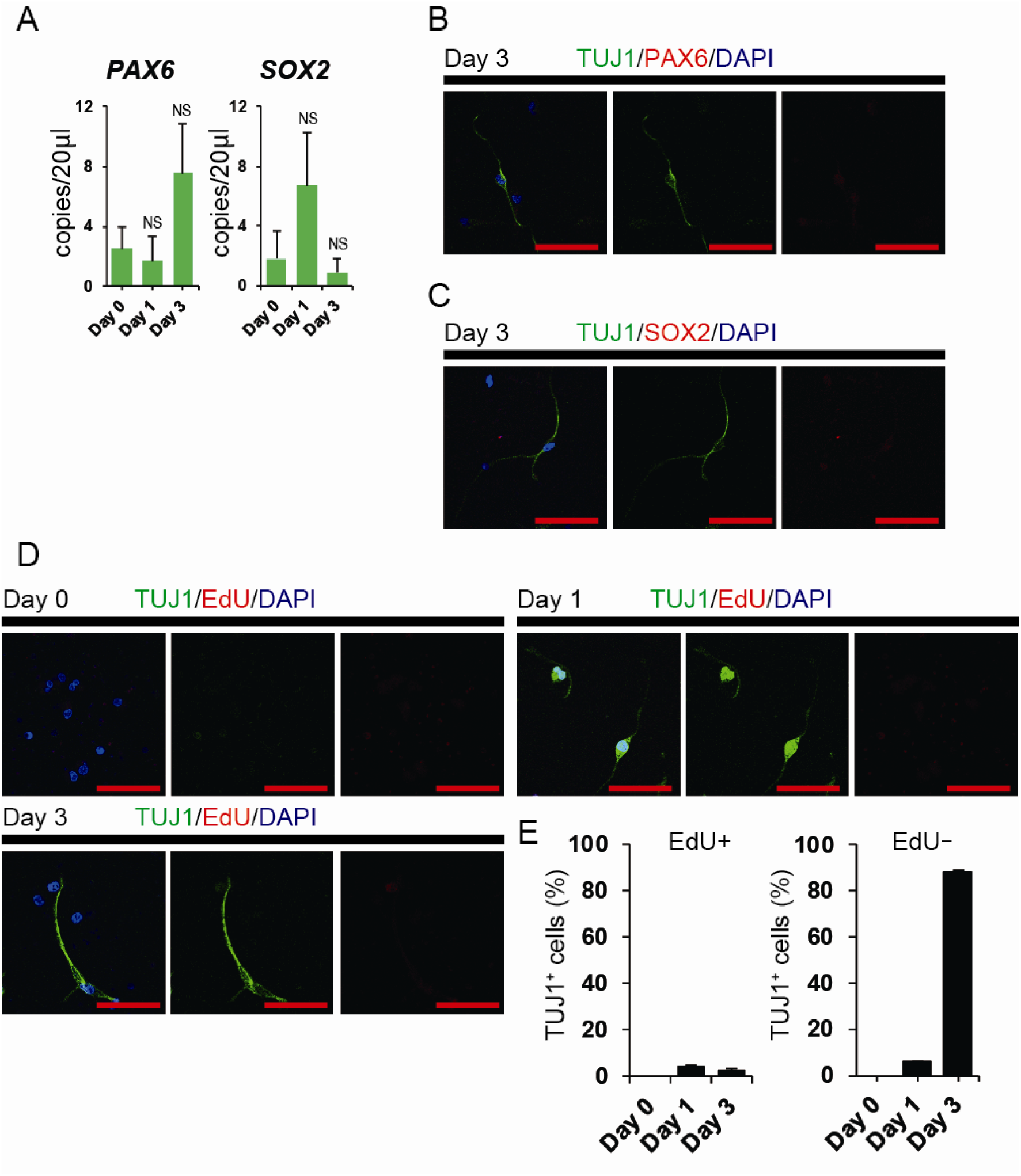
The Blood-to-Neuron Conversion Does Not Involve a Proliferative Neural Progenitor State. (A) mRNA copy number validation of representative neural-progenitor-cells-enriched gene expression using absolute quantification analysis via droplet digital PCR. (mean ± SEM, n = 3 independent experiments; NS, not significant; versus day 0 sample by unpaired t-test). (B-C) Representative immunofluorescence staining for TUJ1 (green), PAX6 (red) and Sox2 (red) at day 3. Scale bar: 50 μm. (E) Representative immunofluorescence staining for EdU (red) at days 0, 1, and 3. Scale bar: 50 μm. (F) Percentage of TUJ1+ cells or EdU- cells among EdU+ cells over the course of reprogramming (day 0 to 3). (mean ± SEM, n = 3 independent experiments)

## Discussion

Herein we report that human adult monocytes may be converted into neuronal cells using a set of small molecules (CDFRVY). While the combination of small molecules, CHIR99021, Dorsomorphin, Forskolin, Repsox, Valproic acid, and Y27632 (CDFRVY) used in this study is a novel combination, each of these molecules has previously been reported to facilitate direct neural conversion in some studies; CHIR99021 and Repsox—the inhibitors of glycogen synthase kinase 3b and transforming growth factor b, respectively—were thought to enhance TFs-mediated neuronal conversion efficiency (Ladewig et al., 2012) and were essential for chemical-mediated neuronal conversion in adult astrocytes (Gao et al., 2017); Dorsomorphine—inhibitor of Bone Morphogenetic Protein signaling pathways (Hong et al 2009)—may be involved in promoting neuronal specification or maturation of iN cells—in collaboration with the TGFβ signaling pathway (Hu et al., 2015); Forskolin reportedly reduced lipid peroxidation, promoted neuronal conversion efficiency (Gascón et al., 2016; Liu et al., 2013) and was important for cell morphology changes (Gao et al., 2017); VPA reportedly promoted neurogenesis, neuronal maturation (Hsieh et al., 2004; Niu et al., 2013) and functioned to activate neuronal genes (Gao et al., 2017); Y27632—selective inhibitor of the rho-associated protein kinase (ROCK) signaling pathways—reportedly facilitated the neuronal conversion of human fibroblasts and resulted in the generation of TUJ1 positive cells with neuron like morphology (Hu et al., 2015). Interestingly, we were unable to convert monocytes into iN cells when using I-BET-151 in addition to CDFRVY—despite reports that I-BET-151 remarkably improved the neuronal conversion efficacy for fibroblasts and astrocytes (Li et al., 2015; Gao et al., 2017)—suggesting that the small molecules required for neural conversion vary depending on the source cell type. Therefore, further investigation is needed to elucidate the complete aspects and roles of these molecules in chemical neuronal conversion.

Our results show that terminally differentiated cells can be transdifferentiated into another, distantly related somatic lineage. The generation of neuronal cells from adult peripheral blood monocytes has important practical implications. Fibroblasts and astrocytes were previously reported to be converted into neuronal cells directly by small molecules (Li et al., 2015, Gao et al., 2017). However, the derivation of fibroblasts and astrocytes requires an invasive and sometimes painful biopsy. By contrast, large numbers of blood monocytes can be obtained via venipuncture. Therefore, the conversion method of monocytes into iN cells detailed in this study allows the generation of human neurons from any individual, unlike the use of fibroblasts and astrocytes as donor cells. In addition, our conversion method does not require the integration of ectopic transgenes and does not pass through the proliferative intermediate state, preventing genomic gene modification and tumorigenesis risks.

From a mechanistic standpoint, we found that, unlike iN cell transdifferentiation from peripheral blood T lymphocytes previously described (Tanabe et al., 2018), feeder cells—like glia cells and fibroblasts—were not required for monocyte transdifferentiation. Although feeder cell systems are useful for laboratory experiments, these support systems contain xenobiotic material (increasing the risk of biological pathogen cross-transfer) and are thus not favorable for future clinical applications.

While this study provides clear evidence that human adult monocytes can be converted into induced neuronal cells expressing neuronal markers, we note that these cells exhibit less mature compared with primary mouse or iPS cell derived iN cells (Matsui et al., 2012; Kim et al., 2011; Maximov et al., 2007; Zhang et al., 2013) according to our result that iN cells consisted mainly of TUJ1/DCX cells (much more than TUJ1/MAP2 cells). The conversion efficacy and neuronal purity of TUJ1/DCX cells were high (around 80-90%); however, monocytes could be converted into TUJ1/MAP2 positive cells with only about 10% efficacy and purity. These results suggest that, although almost all of the monocytes could be coverted to neuronal cell fate initially, the converted cells are immature. Therefore, future efforts should concentrate on improving neuronal maturation, deriving neuronal subtypes, and investigating electrophysiological properties.

## Acknowledgments

We thank Dr. Taisuke Kato of Niigata University for technical support. We also thank all members of the consulted labs for sharing reagents and advice. We would like to thank Editage (www.editage.com) for English language editing. This work was supported by a grant from the Tsubaki Memorial Foundation (Dr Ninomiya).

## Supplementary Information

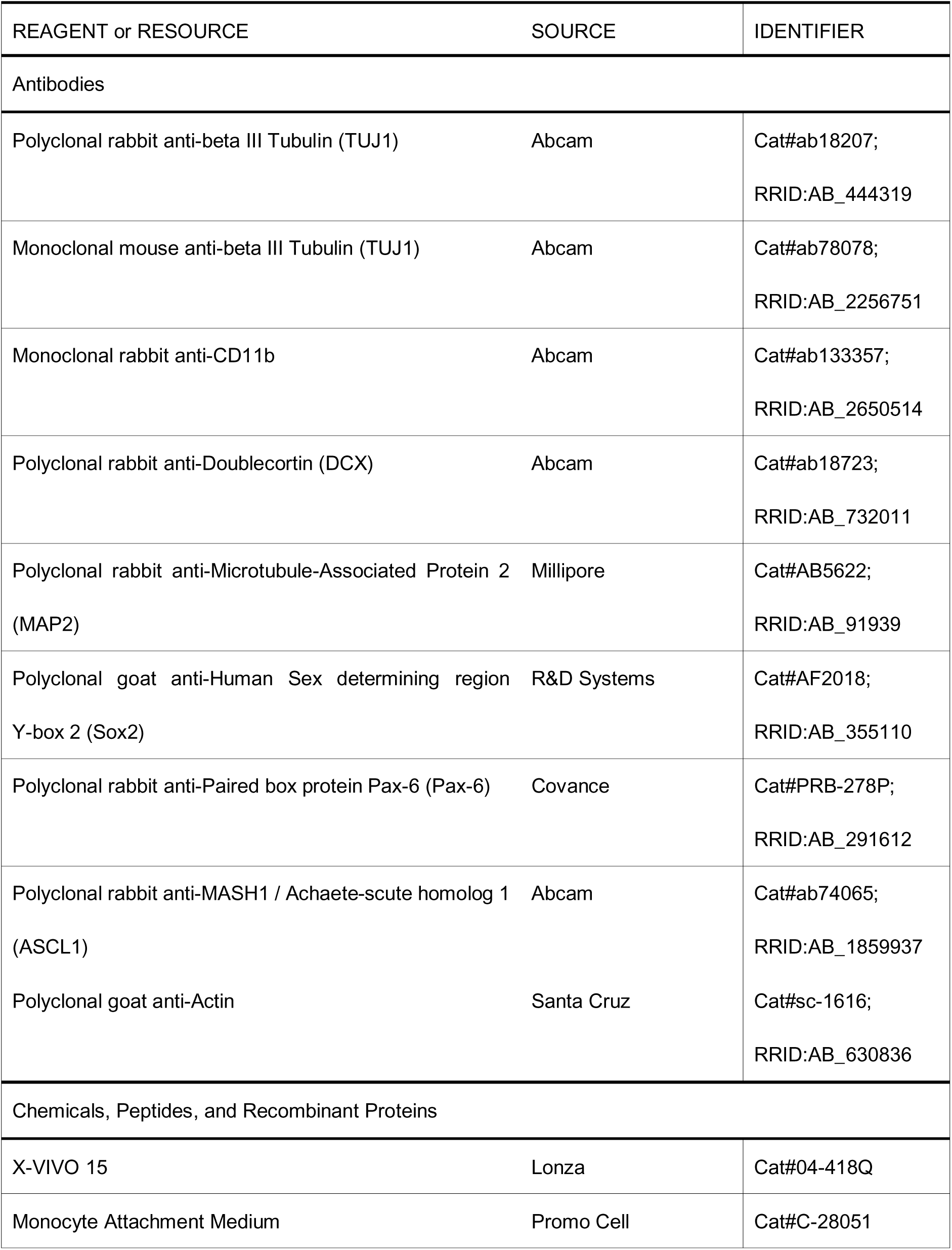
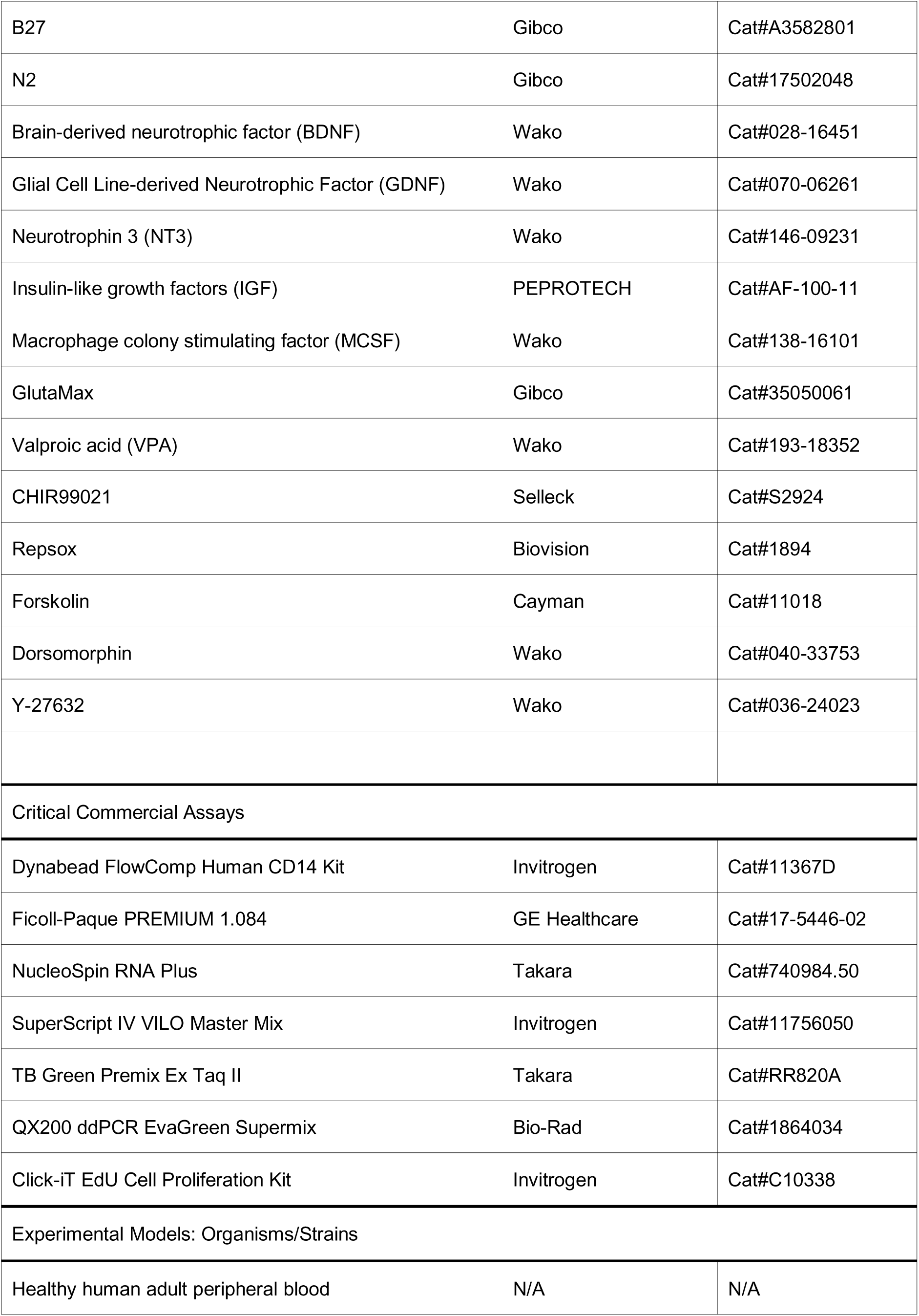
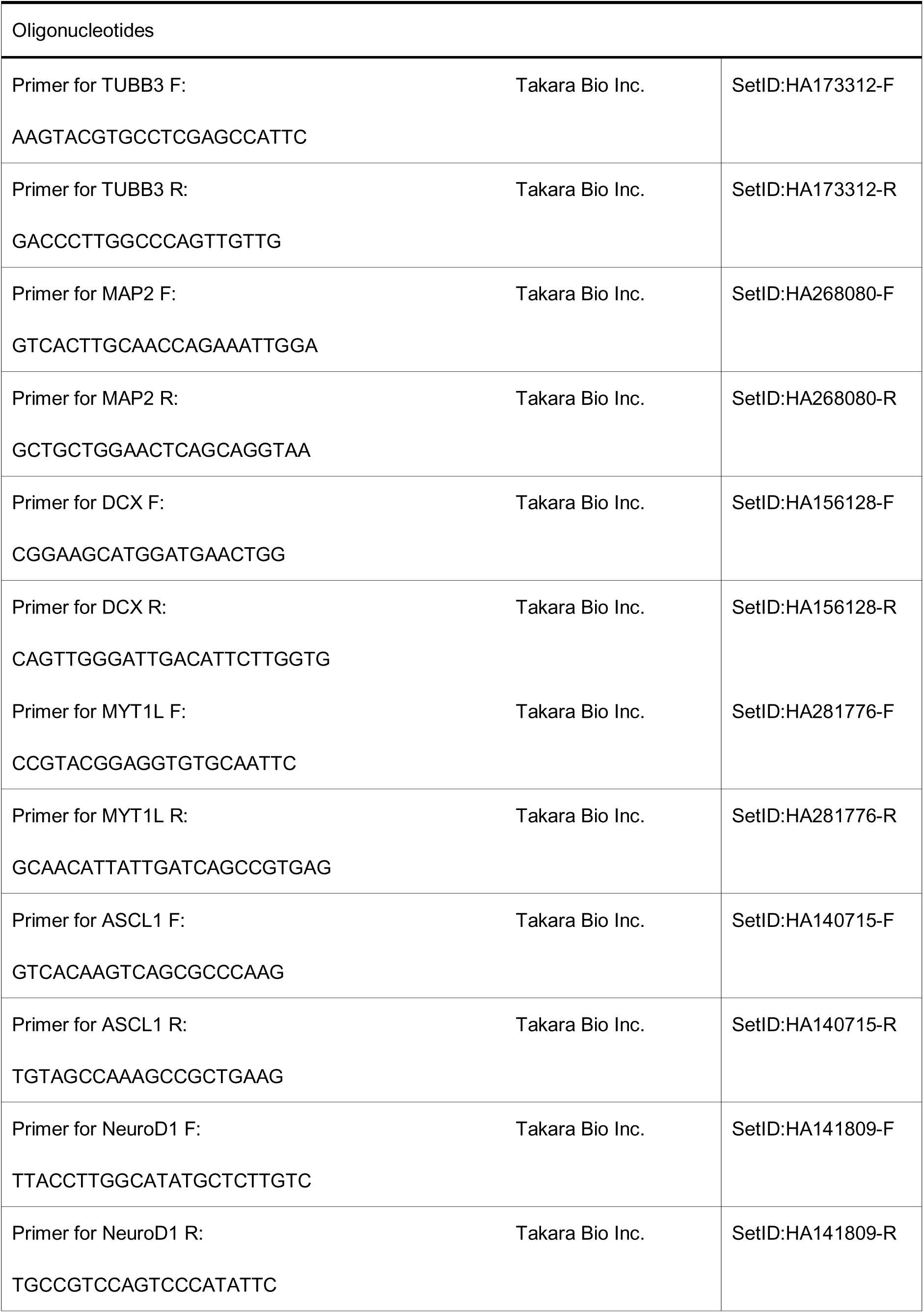
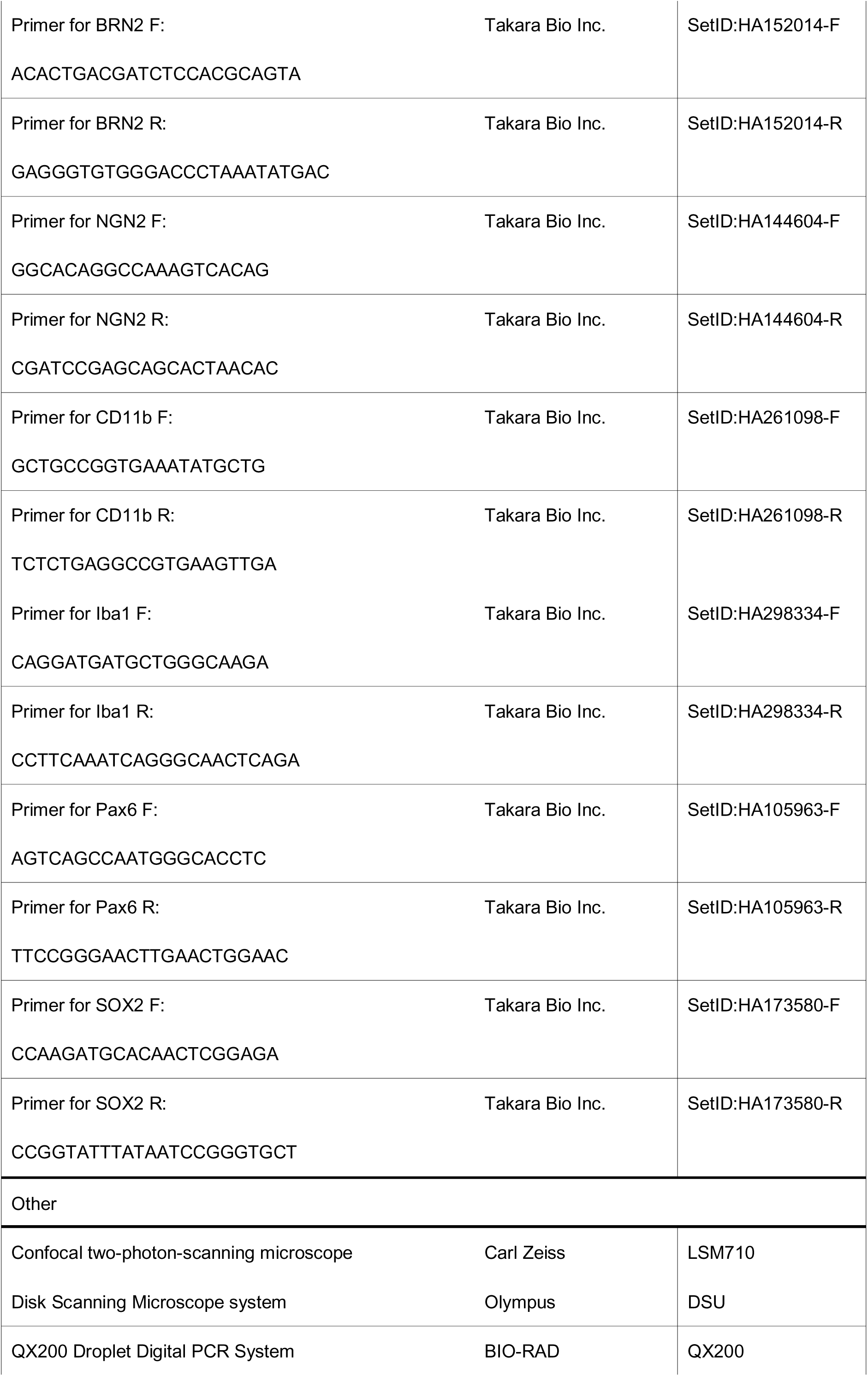

### LEAD CONTACT

Further information and requests for resources and reagents should be directed to and will be fulfilled by the Lead Contact, Itaru Ninomiya

### METHOD

The ethical approval for the present study (#2019-0143) was provided by the Institutional Ethics Committee of the Niigata University Medical and Dental Hospital.

#### Isolation of PBMCs and monocytes

Blood samples were obtained from 8 healthy adult subjects with written consent as approved by the ethics committees of Niigata University Medical and Dental Hospital. Human PBMCs were obtained using the Ficoll-Paque centrifugation (GE Healthcare), according to the manufacturer’s instructions. Primary monocytes and monocyte-depleted PBMCs were isolated from the PBMCs using Dynabeads FlowComp Human CD14 Kit (Invitrogen) according to the manufacturer’s instructions. Cells (1.0 × 10^6^ /cm^2^) were seeded on a poly-D-lysine pre-treated coverslips well in either X-VIVO 15 medium (Lonza) or Monocyte Attachment Medium (MAM) (PromoCell) and incubated at 37 °C for 2 hours. After incubation, culture medium (containing floating cells) was discarded. The remaining adhering cells were washed with fresh PBS and then cultured in conversion medium (either with or without small molecules), defining this time point as day 0.

#### Induction of neuronal cells from human adult monocytes

To generate iN cells, cells were cultured in a conversion medium— consisting of X-VIVO 15 (Lonza) supplemented with 1x B27 (Gibco), 1x N2 (Gibco), 1% GlutaMAX (Gibco), 20 ng/ml BDNF (Wako), 20 ng/ml GDNF (Wako), 10 ng/ml NT3 (Wako), 20 ng/ml IGF (Wako), and 50 ng/ml recombinant human M-CSF (Wako). The conversion medium containing small molecules contained the following small molecules in addition: 0.5 mM VPA (Wako), 3 μM CHIR99021 (Selleck), 1 μM Repsox (Biovision), 10 μ forskolin (Cayman), 2 μ Dorsomorphin (Wako), and 10 µM Y-27632 (Wako) and was changed on day 2.

#### Immunofluorescence staining

Cells plated on glass coverslips were fixed by 4% PFA solution for 5 min at room temperature. After fixation, cells were incubated in 0.3% Triton X-100 for 15 min at room temperature (RT). Cells were then incubated in blocking buffer (1% bovine serum albumin in PBS) for 1 h at RT. Afterward, samples were incubated with primary antibodies at 4 °C overnight. The primary antibodies used were: TUJ1 (1:200, rabbit pAb, Abcam); TUJ1 (1:200, mouse mAb, Abcam); CD11b (1:200, rabbit mAb, Abcam); DCX (1:200, rabbit pAb, abcam); MAP2 (1:500, rabbit pAb, Millipore); Sox2 (1:50, goat pAb, R&D); Pax6 (1:500, rabbit pAb, Covance). The next day, washes were performed with PBS, and samples were incubated with appropriate AlexaFluor secondary antibodies against the appropriate species at 1:1000 (Life Technologies) for 1 h at RT. Nuclei were counterstained with diamino-2-phenylindole, dihydrochloride DNA fluorescent stain (DAPI). Images were captured with a confocal two-photon-scanning microscope (LSM710; Carl Zeiss, Oberkochen, Germany).

Conversion efficiency and neuronal purity were calculated as previously described. Briefly, 10 view fields (×200) were randomly selected for each sample at the indicated time. The number of neuronal cells (TUJ1+/DCX+ or TUJ1+/MAP2+ positive cells with typical neuronal morphology) was counted on day 3 (n = 3). The conversion efficiency was calculated by the ratio of the number of neuronal cells on day 3 to that of initial CD11b+ cells counted in randomly selected 10 view fields on day 0 (n = 3). And the neuronal purity was calculated by the ratio of the number of neuronal cells to the total cell number indicated by DAPI.

#### Western blotting

For the *in vitro* whole-cell extracts, cells were harvested using RIPA Lysis Buffer System (Santa Cruz, sc-24948). Proteins in the samples were separated by tris-glycine SDS-PAGE and probed with primary antibodies. The following primary antibodies were used: TUJ1 (1:1000, mouse mAb, Abcam); CD11b (1:1000, rabbit mAb, Abcam); DCX (1:1000, rabbit pAb, Abcam); Ascl1 (1:2000, Rabbit pAb, Abcam); Actin (1:10000, goat pAb, Santa Cruz). Detection was achieved using Horse Radish Peroxidase (HRP)-linked secondary antibodies (against the appropriate species) and ECL Plus Western Blotting Detection System (GE Healthcare). Densitometry was performed using ImageJ software. Protein levels were normalized to the Actin loading control.

#### Quantitative real-time PCR and Droplet digital PCR

The total RNA of indicated cell samples was isolated using NucleoSpin RNA Plus (Takara) following manufacturer’s instructions. This RNA was used for reverse transcription with SuperScript IV VILO Master Mix (Invitrogen) according to manufacturer’s instructions. To select normalization factors, we calculated the relative stability of 16 housekeeping genes’ expression (Human Housekeeping Gene Primer Set, Takarabio) in each cultured cell using the free analysis program RefFinder (heartcure.com.au/reffinder/). Two genes (RPLP2 and RPS18) were selected as stable endogenous expression controls. Quantitative real-time PCR was conducted with 400 nM primers, 25 ng cDNA template and TB Green Premix Ex Tax II (Takara, RR820A) in Thermal Cycler Dice Real Time System (Takara, Software Ver. 5.11B for TP800) (n = 3).

Droplet digital PCR was performed on the Q×200 droplet digital PCR system (Bio-Rad). Briefly, Taq polymerase PCR reaction mixtures were assembled using 2× QX200 ddPCR EvaGreen Supermix, 25 ng cDNA template and 250 nM primers. DG8 cartridges were loaded with 20 μL PCR reaction mixtures and 70 μL of droplet generation oil for each sample. The cartridges were placed into a droplet generator for emulsification and the emulsified samples were transferred onto 96-well plate as per manufacturer’s instructions. Plates were heat-sealed and underwent 40 cycles in a simpliAmp thermal cycler (ThermoFisher). After PCR, the 96-well droplet PCR plates were loaded into a droplet reader, which sequentially reads the droplets from each well of the plate. Absolute quantification analysis of the ddPCR data was performed using QuantaSoft (version 1.7.4.0917) and the mRNA copy number of each target gene was calculated using RPLP2 and RPS18 as reference genes. All primers were bought from Takara Bio Inc.

#### EdU incorporation assay

EdU incorporation and EdU staining were performed using a Click-it EdU Cell Proliferation Kit for Imaging (Invitrogen) according to manufacturer’s instructions with slight modification. Briefly, 10 μ EdU was added 2 h prior to fixation on indicated days. Cells were fixed by 4% PFA for 5 min and permeabilized with 0.3% Triton X-100 in PBS at room temperature. The EdU staining was performed following the manufacturer’s instructions. Images were captured and analyzed with a confocal two-photon-scanning microscope (LSM710; Carl Zeiss, Oberkochen, Germany).

### QUANTIFICATION AND STATISTICAL ANALYSIS

All quantified data were statistically analyzed and are presented as mean ± SEM. Differences in the parameters were evaluated using the unpaired t-test. All statistical analyses were performed using BellCurve for Excel, Version 3.20 (Social Survey Research Information Co., Ltd.).

